# pFLEX – a Python library for fast functional evaluation of genetic networks at the biological module-level

**DOI:** 10.64898/2026.06.11.731557

**Authors:** Talip Yasir Demirtas, Angela Helen Shaw, Maximilian Billmann

## Abstract

Genetic networks derived from omics data are a powerful tool for systematic gene function prediction. Performance evaluation of such predictions is crucial to judge the data and computational pipeline for network construction, but unbalanced functional standards often cause hidden evaluation biases. To visualize and mitigate such biases, we previously developed the R package FLEX. Here, we present the pFLEX genetic network benchmarking tool as Python library with new and improved functionality. pFLEX improves overall runtime 4.1 to 15.8-fold. It offers additional evaluation metrics that allow for easy comparison of precision recall performance at the complex or pathway resolution between genetic networks. We demonstrate the utility of pFLEX for evaluating tissue-specific co-essentiality networks and data normalization strategies of the Cancer Dependency Map, as well as for cell line-specific Perturb-Seq-derived networks. This illustrates the requirement for biological module-resolved precision recall metrics in pFLEX for sensitive and fast evaluation of genetic networks.

**Availability and Implementation:** pFLEX is available under the MIT license at https://github.com/billmannlab/pFLEX and the pFLEX version used in this manuscript along with benchmarking code for the analyses presented in this manuscript are archived at https://doi.org/10.5281/zenodo.20632868.

## Introduction

Genetic networks are a powerful tool for predicting gene function. In the network, genes are represented as nodes connected by edges that represent their functional relation (Billmann et al. 2026). Those functional relations can illustrate that proteins act in the same complex or pathway or mediate the response to environmental changes. Functional relations are quantitative, allowing both the simple classification of genes into modules as well as drawing a more complex wiring diagram of the cell. While genetic networks can be constructed using various omics data, gene perturbation data, which records cellular phenotypes upon the systematic inactivation or activation of genes has become particularly powerful (Wainberg et al. 2021, Billmann et al. 2026).

In human cells, the precise perturbation of thousands of genes in a single experiment has been made possible through CRISPR-based gene editing tools. To derive genetic networks from genetic perturbation data multiple features for each gene are recorded. Perhaps the biggest strength of those networks is that relatively general measurements such as cell fitness allow for reconstruction of highly specific functional modules of the cell, rendering networks an unbiased, universal discovery tool for the genotype to phenotype problem (Wainberg et al. 2021, Billmann et al. 2026).

To extract accurate measurements from CRISPR screens, processing and normalization of the read-count-based raw data is crucial. Sophisticated normalization tools have been developed not only to account for technical artifacts but also for false positives due to the biology of the experimental system (Meyers et al. 2017, Iorio et al. 2018, Allen et al. 2019, Dempster et al. 2021, Gheorghe and Hart 2022, Hassan et al. 2023, Billmann et al. 2025). Importantly, global performance metrics such as precision-recall (PR) analysis that treat all co-annotated gene pairs the same, often fail to visualize and quantify the success of such normalization techniques. To address this, we recently developed FLEX, an R package to systematically benchmark CRISPR screen-based network performance at the biological module level (Rahman et al. 2021).

Here, we present pFLEX, a Python library for fast functional evaluation of genetic networks at the biological module-level. pFLEX benchmarks network up to fifteen times faster than the R implementation. pFLEX provides an additional per-module data set comparison function. To illustrate its utility, we show how CRISPR screens performed in cell lines derived from different tissue types reconstruct different biological processes and how normalizing for covariance patterns in the Cancer Dependency Map (DepMap) increases the diversity of captured biological processes without affecting global PR performance. Finally, we resolve biological complex-specific gene embeddings in Perturb-Seq data from different cell lines.

## Results and discussion

pFLEX includes all network evaluation functions implemented in FLEX/R and added new functions to enable better comparability of networks. pFLEX measures global performance of a network to assign gene pairs to modules (complexes, pathways, biological processes) of a given functional standard and visualizes how each module contributes to the PR curve via the contribution diversity plot (Figure 1A). pFLEX offers a module-level (m)PR curve that treats recovered modules (instead of gene pairs) as true positives, which balances the global PR metric accounting for unequal module size. At the module-level, pFLEX compares area under the PR curve (AUPRC) with the size of a module and now also allows to compare per-module AUPRC between two networks. Finally, pFLEX now counts the number of modules passing a given AUPRC threshold to compare networks with a simple summary statistic.

**Figure 1.**
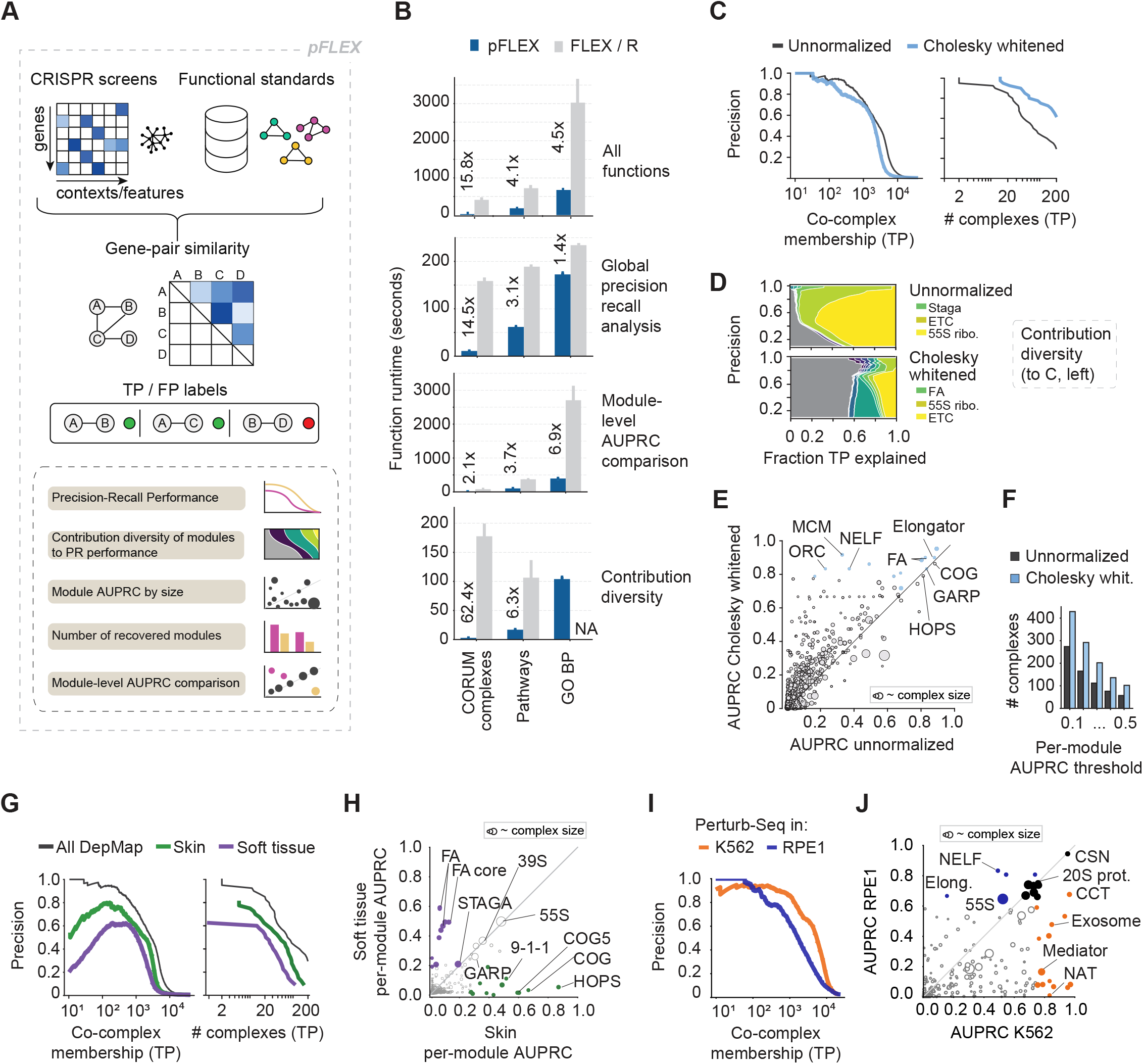
pFLEX workflow, runtime performance, and module-resolved benchmarking of bulk and single cell CRISPR screening-derived genetic networks. **(A)** Schematic overview of the pFLEX workflow. Gene-effect matrices from CRISPR screens and curated functional standards are used to compute ranked gene-pair similarities, assign true-positive (TP) and false-positive labels based on functional co-membership, and generate global precision-recall and module-level performance outputs. **(B)** Runtime comparison between pFLEX and FLEX/R across CORUM complex, pathway, and GO BP functional standards. Bars show mean wall-clock runtime across 10 runs for the full workflow and selected major analysis steps. Error bars show standard deviations of the 10 runs. Labels indicate the fold speedup of pFLEX over FLEX/R. **(C)** Global (left) and module-level (right) precision-recall (PR) performance comparison of 25Q2 DepMap-derived networks before and after Cholesky whitening. **(D)** Contribution diversity in global PR curves of unnormalized and Cholesky whitened networks. If most of the area, especially at higher precision, is dominated by the colored top10 modules (of approx. 2000), contribution diversity is low and PR performance dominated by few biological modules. **(E)** Comparison of per-complex AUPRC values between Cholesky whitened and unnormalized networks. **(F)** Number of modules (here complexes) with per-module AUPRC performance matching the indicated threshold. This summarizes the numbers also plotted in panels C (right) and E. **(G)** Global (left) and module-level (right) PR performance of DepMap-derived co-essentiality networks constructed from all 1183 25Q2 DepMap genome-wide CRISPR screens, skin (n = 75) and soft-tissue (n = 46) -derived cell lines. Precision is shown as a function of recovered CORUM co-complex memberships (TP). Module-level PR analysis summarizes how many functional modules (here complexes) are recovered at different precision levels. **(H)** Comparison of per-complex AUPRC values between skin-derived and soft-tissue-derived networks. Labeled complexes highlight functional modules that are better recovered in one tissue-specific network. **(I)** Global PR performance comparison of genetic networks constructed from Perturb-Seq data targeting essential genes in K562 and RPE1 cells. Expression profiles are used as feature profiles to estimate pairwise gene similarity. **(J)** Comparison of per-complex AUPRC values between K562 and RPE1 Perturb-Seq-derived networks.

The implementation of those evaluation functions in pFLEX now allows for an easy benchmarking of genetic networks in the Python ecosystem. Importantly, this implementation achieved shorter total runtimes across all tested functional standards, with speedups of 15.8-fold for the CORUM protein complex standard, 4.0-fold for pathway annotations, and 4.5-fold for the Gene Ontology Biological Process (GO BP) standard. For protein complex comembership evaluation of a DepMap-derived co-essentiality network, pFLEX completed the benchmark in 26.0 seconds on a Windows 11 Pro workstation equipped with an Intel Core i9-14900K CPU, 24 cores, 32 logical processors, and 128 GB RAM, compared with 409.5 seconds for FLEX/R (Figure 1B).

To illustrate the functionality of pFLEX including its new functions, we benchmarked two sets of networks derived from the 25Q2 DepMap: covariance normalization of the global network and tumor type-specific networks. First, we characterized covariance normalization via Cholesky whitening of a DepMap-derived genetic network. Only mPR curves but not global PR curves, which are routinely used for benchmarking, captured the substantially increased biological diversity in the network after normalization (Figure 1C). The source of this improved performance can be visualized by the contribution diversity function (Figure 1D). Those three functions are also available in FLEX/R (Rahman et al. 2021). However, pFLEX fixed an issue in the contribution diversity function that prevented the FLEX/R version to complete with the largest functional standard GO BP (Figure 1B). When further benchmarking DepMap normalization pFLEX per-complex AUPRC scatter function now enables a direct comparison of how much AUPRC performance of each complex changes following normalization (Figure 1E). pFLEX also now compares the number of complexes that pass a given AUPRC threshold in different networks (Figure 1F).

Second, we utilized pFLEX to compare genetic networks constructed from DepMap cell lines of different origin. We selected two tumor entities: skin with 75 and soft tissue with 46 cell lines. pFLEX first showed how well both DepMap sub-networks as well as the complete DepMap network (1183 cell lines) reconstruct co-complex memberships with and without accounting for complex size (Figure 1G). pFLEX now allows to compare per-complex AUPRC performance between different networks, illustrated by comparing DepMap sub-networks based on skin and soft tissue-derived cell lines (Figure 1H). This showed that the skin-based network reconstructed complexes such as the HOPS or COG complexes, whereas the soft tissue-based network reconstructed DNA damage repair complexes.

Finally, we used pFLEX to compare genetic networks constructed from Perturb-Seq data in K562 and RPE1 cells (Replogle et al. 2022). Those data contain perturbations of essential genes and the pairwise similarity between perturbed genes is estimated by computing the Pearson correlation coefficient along expression profiles measured in single cells. Again, while the global PR curve generally compared network performance, only the pFLEX per-complex AUPRC performance scatter plot showed the many complexes performing differently between both networks (Figure 1I, J)

Together, pFLEX enables quantitative benchmarking of genetic networks at the biological module level in the Python ecosystem. It substantially improves runtimes over the original FLEX/R benchmarking framework and adds additional functions for a more detailed comparison of different networks, demonstrated by the comparisons of genetic networks constructed from the DepMap and Perturb-Seq data.

## Implementation and methods

### Installation of pFLEX

pFLEX can be installed from *PyPI* using “*pip install pflex”*. For local development and reproduction of manuscript analyses, the source version can be installed from GitHub by cloning https://github.com/billmannlab/pFLEX and running “*uv pip install -e* .*”* from the repository root.

### Overview of the pFLEX workflow

pFLEX reimplements the FLEX/R benchmarking framework (Rahman et al. 2021) in Python to address scalability and runtime limitations observed when evaluating genome-wide genetic networks with comprehensive functional standards such as GO BP. pFLEX also streamlined the usability of the different functions to make it more intuitive. The workflow is structured around three core stages: (i) functional standard preparation, which loads and filters curated gene sets to define reference biological modules; (ii) performance evaluation, which ranks gene–gene similarity relationships and quantifies global and module-level performance; and (iii) visualization, which summarizes performance, module-specific metrics and performance-contributing patterns. In addition to FLEX/R functionality, pFLEX now supports cross-dataset comparison. The workflow is summarized in Figure 1A.

### Runtime-oriented implementation of pFLEX

Runtimes primarily improved through optimization of pairwise similarity handling and pre-indexed functional standard lookups, which avoids and reduces both computation and memory pressure repeatedly rebuilding expensive gene-pair mappings during module-level and global precision-recall analysis. Computational bottlenecks present in FLEX/R, particularly when using large functional standards such as GO BP, are reduced through vectorized array operations and compiled routines. Correlation, sorting, and pair filtering run closer to native numerical-code speed (Harris et al. 2020).

### Functional standard preparation

Functional standards in pFLEX, just as in FLEX/R, provide an approximated ground truth for functional relations of genes by reporting co-annotation to protein complex, pathways or processes. Functional standards are loaded either from built-in resources, including CORUM 3.0 protein complexes (Giurgiu et al. 2019), GO BP annotations (Ashburner et al. 2000), and curated pathway collections (Liberzon et al. 2011), or from user-supplied annotations. Redundant complexes are removed by excluding entries with identical gene membership. To focus on specific biological functions, users may remove specific modules from the functional standard by name or by size. For the selected functional standard, pFLEX constructs an internal representation of gene–gene relationships. Gene pairs co-annotated within the same functional unit are labeled as positives (1), whereas non–co-annotated pairs are labeled as negatives (0). In pFLEX, functional standards are processed once at initialization, harmonized to gene-level identifiers, and intersected with each dataset. All downstream analyses are restricted to genes present in the standard.

### Input data and similarity ranking

pFLEX benchmarks genetic networks (i.e. genes connected by edges representing similar feature profiles) constructed from gene perturbation datasets. As in FLEX/R, two input formats are accepted. First, the input data are arrays of perturbated genes as rows features as columns. Features are experimental backgrounds (cell lines, mutations) when cell fitness is measured or expression profiles from e.g. Perturb-Seq experiments. From gene-by-feature matrices, pairwise gene similarity scores are estimated by computing Pearson correlation coefficients across features. Second, provided similarity matrices are directly used.

For this study, gene perturbation effect scores from the Cancer Dependency Map (DepMap) version 25Q2 were used. Those comprised 1183 genome-wide CRISPR-Cas9 screens in cultured cell lines targeting 17,916 genes. Moreover, Perturb-Seq data targeting essential genes in K562 (2057 perturbed genes) and RPE1 (2393 perturbed genes) cells, restricted to the most variably expressed 2320 (K562) and 2111 (RPE1) genes on the feature side, were used (Replogle et al. 2022).

### Global precision-recall benchmarking

As FLEX/R, pFLEX quantifies the performance of genetic networks to recover known functional gene relations by comparing ranked gene–gene similarities and functional standards, summarized using precision-recall (PR) metrics, which are robust results to class imbalance (Myers et al. 2006, Saito and Rehmsmeier 2015). Rather than plotting recall directly, pFLEX displays precision as a function of the cumulative number of true positives (TPs) along the ranked list. The resulting precision–TP curve shows how quickly true functional relationships accumulate as ranking thresholds relax, and its area (area under the PR curve; AUPRC) summarizes global functional performance.

### Module-level benchmarking

As FLEX/R, pFLEX evaluates performance at the level of individual functional modules (complex, pathway or process). While the global PR analysis ranks all gene pairs once and computes a single precision–TP curve for a complete functional standard, the per-module analysis assesses each module independently using the same ranked gene–gene similarity list. Co-annotated gene pairs for a given module are treated as positives for that module, while other pairs represent negatives. Precision and cumulative true positives are recomputed within this restricted subset, and an AUPRC is computed for each module separately (instead of a single global metric). Per-module AUPRC performance can be contrasted to complex size to identify biases in global PR performance associated with large modules. pFLEX now allows to contrast module-level AUPRC values from different networks.

### Module-level recovery summary

Beyond per-module AUPRC scores, pFLEX (just as FLEX/R) provides a module-level summary that quantifies recovered modules (instead of gene pairs). Using the globally ranked gene-gene similarities, true positives are assigned to modules in a stepwise manner along increasing precision, estimating the number of distinct recovered complexes under defined coverage criteria (Rahman et al. 2021). This results in the module (m)PR curve, a metric that mitigates evaluation bias. Contribution diversity plots show the fraction of TP gene pairs that belong to distinct modules along precision thresholds, visualizing evaluation bias in global PR curves.

### Visualization and exported outputs

pFLEX provides visualization outputs aligned with the three evaluation stages described above. For global performance (Performance evaluation I), precision-TP curves display how true positives accumulate along the ranked gene–gene similarity list, with AUPRC values reported as a compact dataset-level metric. For module-level performance (Performance evaluation II), per-module AUPRC scatter plots highlight heterogeneity in performance across biological neighbourhoods. High-AUPRC modules can be highlighted and per-module AUPRC can be plotted against complex size. In multi-dataset settings, module-level metrics can be compared across datasets. For module-level summary (Performance evaluation III), mPR curves are plotted. pFLEX also quantifies contribution diversity in global PR performance using a greedy approximation algorithm that allocates true positives stepwise across modules along precision levels.

pFLEX visualization outputs are accompanied by exported tables to facilitate downstream analyses and reproducibility.

### Cross-dataset comparison

In pFLEX, multiple datasets can be evaluated within a single run using the same functional standards and ranking procedures. Differences between datasets can be examined both at the global level and at the module level using comparative PR and module-level AUPRC metrics.

### Normalization comparison workflow

To benchmark data normalization effects in genetic networks, we applied Cholesky whitening to the DepMap gene effect matrix before computing Pearson’s correlation coefficients as pairwise similarity scores. This transformation addressed correlated structures across samples by rescaling the gene-by-sample covariance matrix to the identity (Gheorghe and Hart 2022). The Cholesky whitened network was benchmarked against the network derived from the unnormalized data.

## Data availability

No new experimental data were generated in this study. pFLEX is available under the MIT license at https://github.com/billmannlab/pFLEX. The pFLEX library contains mock data comprising scores for genes perturbed in the 25Q2 DepMap that overlap with the CORUM 3.0 functional standard in skin and soft tissue-derived cell lines (also shown in Figure 1G, H). The pFLEX version used in this manuscript along with benchmarking code enabling reproducibility of analyses presented in this manuscript are archived at https://doi.org/10.5281/zenodo.20632868.

## Acknowledgements

We thank all members of the Billmann lab for fruitful discussions.

## Author contributions

T.Y.D. conceived the project, implemented the software, performed analyses and drafted the manuscript. M.B. conceived the project, interpreted analyses and wrote the manuscript. A.H.S. contributed to analyses and tested the software.

## Funding

M.B. is partially funded by the DFG TRR259.

## Conflicts of interest

None declared.

## Notes

https://github.com/billmannlab/pFLEX

### Competing Interest Statement

The authors have declared no competing interest.

